# Serine Palmitoyltransferase Controls Stemness of Intestinal Progenitors

**DOI:** 10.1101/2020.12.03.409128

**Authors:** Ying Li, Bhagirath Chaurasia, Vincent Kaddai, Joseph L. Wilkerson, J. Alan Maschek, James Cox, Peng Wei, Claire Bensard, Peter J Meikle, Hans Clevers, James A Shayman, Yoshio Hirabayashi, William L. Holland, Jared Rutter, Scott A. Summers

## Abstract

Cancers of the gastrointestinal tract including esophageal adenocarcinomas, colorectal cancers, and cancers of the gastric cardia are common comorbidities of obesity. Excessive delivery of macronutrients to the cells lining the gut can increase one’s risk for these cancer by inducing imbalances in the rate of intestinal stem cell proliferation vs. differentiation, which can produce polyps and other aberrant growths. We demonstrate that serine palmitoyltransferase (SPT), which diverts dietary fatty and amino acids into the sphingolipid biosynthesis pathway, is a critical modulator of intestinal stem cell homeostasis. SPT and other enzymes in the biosynthetic pathway are upregulated in human colon tumors. These enzymes produce sphingolipids that serve as pro-stemness signals that stimulate peroxisome-proliferator activated receptor alpha (PPARα)-mediated induction of fatty acid binding protein-1. This increases fatty acid uptake and oxidation and enhances the stemness program. Serine palmitoyltransferase thus serves as a critical link between dietary macronutrients, epithelial regeneration, and cancer risk.

## Introduction

The intestinal epithelium is the most rapidly regenerating tissue in the body. Constant mechanical damage inflicted by passing bowel content necessitates the nearly complete renewal of the epithelium every 4–5 days (Leblond and Stevens, 1948). Replacement of the intestinal villi results from a coordinated process triggered by the proliferation and differentiation of the intestinal stem cells (ISCs) that lie in a crypt at the base of the villus. The ISCs give rise to the specialized absorptive cells (i.e. enterocytes) and mucus and hormone-secreting goblet and enteroendocrine cells, respectively, that comprise the bulk of the villus epithelium. The balance between ISC proliferation and differentiation is tightly regulated and its disruption leads to gastroenteritis or hyperproliferative lesions and tumors.

ISCs respond to nutritional cues that increase proliferation and enhance the regenerative capacity of the gut epithelium. For example, the fatty acid palmitate stimulates proliferation of ISCs, thus increasing the number of cells within the crypt to enhance tumor risk (Beyaz et al., 2016). By contrast, serine deprivation restricts growth of intestinal cancers (Maddocks et al., 2017). Since palmitate and serine are required to synthesize the sphingoid backbone of sphingolipids, we developed the hypothesis tested herein that one or more sphingolipids might be important signaling intermediates linking these dietary macronutrients to stem cell fate.

Once palmitate enters cells, it is quickly conjugated to coenzyme-A, which traps the fatty acid in the cell and activates it for subsequent metabolism. The resultant palmitoyl-CoA can be coupled to (i) a glycerol backbone to produce glycerophospholipids, (ii) carnitine for transport into mitochondria, or (iii) the aforementioned amino acid serine to generate sphingolipids. This latter serine conjugation step is catalyzed by serine palmitoyltransferase (SPT), which generates the sphingoid backbone present in sphingolipids. This moiety subsequently acquires additional, variable fatty acids and a critical double bond to form ceramides, which are the precursors of sphingomyelins and gangliosides (Figure 1a). Ceramides, particularly those containing a C16-acyl chain, have emerged as important nutrient signals that alter cellular metabolism in response to excessive ectopic fatty acids (Summers et al., 2019). Though sphingolipids are relatively minor components (Figure 1B, light blue) of the mouse gut lipidome that are far less abundant than glycerolipids and sterols (Figure 1B, dark blue), the studies presented herein reveal that they are potent nutrient signals that alter metabolism of ISCs to enhance stemness and catalyze epithelial regeneration.

**Figure 1.**
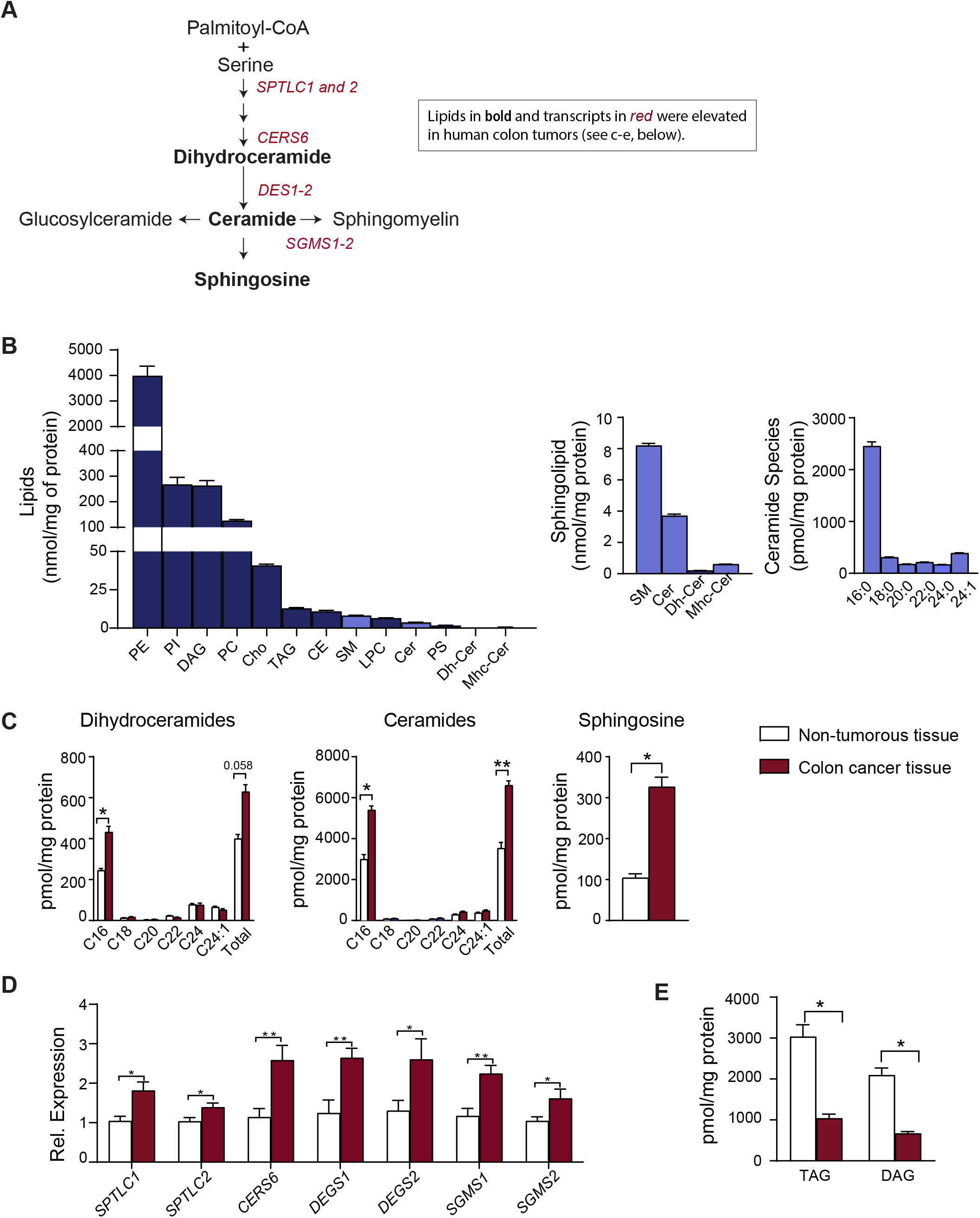
Sphingolipids and sphingolipid-synthesizing genes are upregulated in human colon cancer. (A) Rough schematic of the sphingolipid synthesis pathway. Shown in bold font are the lipids, and in red the transcripts, that were upregulated in human colon cancer (see C and D, below). (B) Major lipid species in the mouse small intestine. Sphingolipids are in light blue. (C) Quantification of sphingolipids in colon cancer tissue and adjacent non-tumorous tissue from patients. (D) qPCR analysis of transcripts encoding sphingolipid synthesizing enzymes in colon cancer tissue or adjacent non-tumorous tissue obtained from patients. (E) Triacylglycerol and diacylglycerol in colon cancer tissue and adjacent non-tumorous tissue obtained from patients. (*p<0.05, **p<0.01, ***p<0.001, n=9). Abbreviations: PE, phosphatidylethanolamine; PI, phosphatidylinositol; DAG, diacylglycerol; PC, phosphatidylcholine; Cho, cholesterol; TAG, triacylglycerol; CE, cholesterol esthers; SM, sphingomyelin; LPC, lysophosphatidylcholine; Cer, ceramide; PS, phosphatidylserine; Dh-Cer, dihydroceramide; Mhc-Cer, monohexosylceramide; *Sptlc1*, serine palmitoyltransferase long chain base subunit 1; Sptlc2, serine palmitoyltransferase, long chain base subunit 2; *Cers6*, ceramide synthase-6; *Degs1*, dihydroceramide desaturase-1; *Degs2*, dihydroceramide desaturase-2; *Sgms1*, sphingomyelin synthase-1; *Sgms2*, sphingomyelin synthase-2.

## Results

### Sphingolipids and sphingolipid-synthesizing enzymes are upregulated in human colon cancer

Adenocarcinomas that were surgically removed from colon cancer patients (Table 1) contained higher levels of C_16_-ceramides, C_16_-dihydroceramides, and sphingosine than the healthy tissue surrounding the lesion(Figure 1C). They also contained higher levels of transcripts encoding two critical SPT subunits (*SPTLC1&2)* and several other sphingolipid-synthesizing enzymes (i.e. *CERS6*, *DEGS1&2*, and *SGMS1&2*) (Figures 1D). *CERS6* encodes the ceramide synthase that is required to make the C_16_-ceramides that are signals of nutritional excess (Hammerschmidt et al., 2019; Raichur et al., 2019; Raichur et al., 2014; Turpin et al., 2014) and are elevated in the human tumors (Figure 1C). By comparison, the other *CERS* isoforms (1-5), and the ceramides they produce, were unaltered in the lesions (Figure 1C and S1B). The *SGMS* transcripts that encode the sphingomyelin synthases that convert ceramide into sphingomyelin were slightly elevated, but sphingomyelins were not (Figure S1A). Moreover, other genes that influence ceramide synthesis or metabolism were unaltered (Figure S1B). Thus, the increase in transcripts and sphingolipids was largely restricted to the enzymes in the *de novo* synthesis pathway that produce the C_16_-ceramides. We also quantified two glycerol-containing lipids (i.e. diacylglycerol and triacylglycerol), finding that they were reduced in the tumors (Figure 1E). This finding suggests that these intestinal tumors redirect fatty acids away from the glycerolipid pathway and towards sphingolipids. As expected, the tumors contained substantially higher levels of *LGR5*, a marker of ISCs (Figure S1C).

**Table 1:**
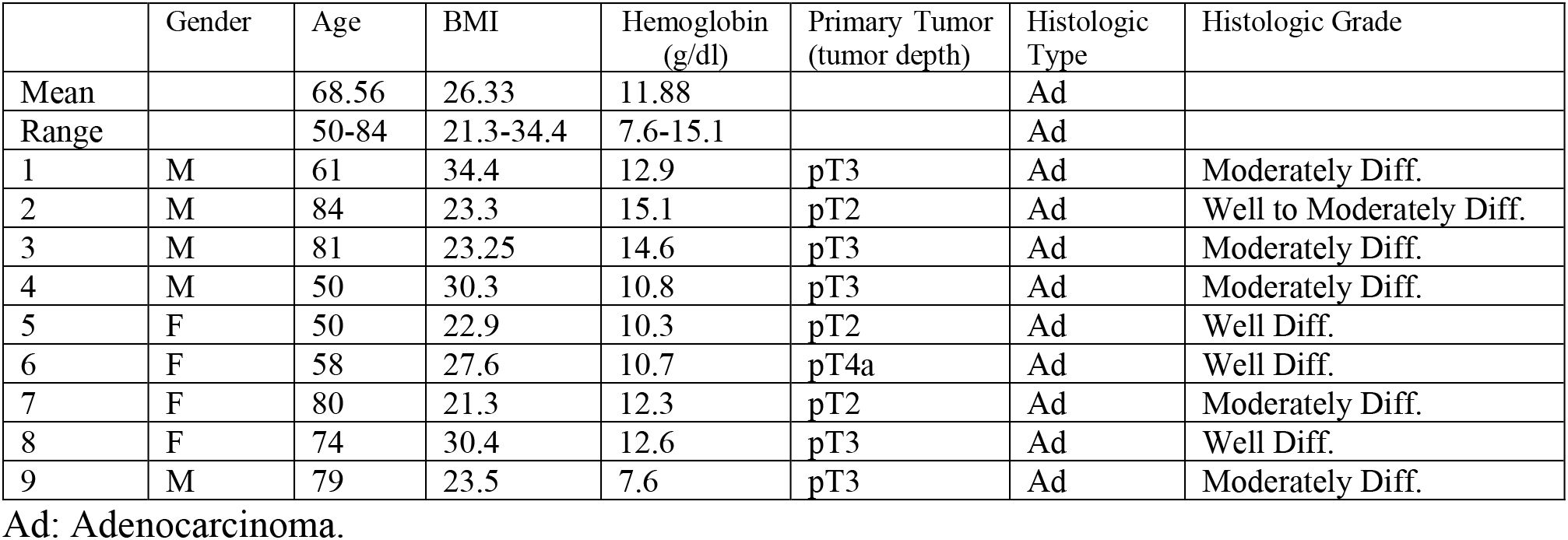
Patient information

|  | Gender | Age | BMI | Hemoglobin (g/dl) | Primary Tumor (tumor depth) | Histologic Type | Histologic Grade |
| --- | --- | --- | --- | --- | --- | --- | --- |
| Mean |  | 68.56 | 26.33 | 11.88 |  | Ad |  |
| Range |  | 50-84 | 21.3-34.4 | 7.6-15.1 |  | Ad |  |
| 1 | M | 61 | 34.4 | 12.9 | pT3 | Ad | Moderately Diff. |
| 2 | M | 84 | 23.3 | 15.1 | pT2 | Ad | Well to Moderately Diff. |
| 3 | M | 81 | 23.25 | 14.6 | pT3 | Ad | Moderately Diff. |
| 4 | M | 50 | 30.3 | 10.8 | pT3 | Ad | Moderately Diff. |
| 5 | F | 50 | 22.9 | 10.3 | pT2 | Ad | Well Diff. |
| 6 | F | 58 | 27.6 | 10.7 | pT4a | Ad | Well Diff. |
| 7 | F | 80 | 21.3 | 12.3 | pT2 | Ad | Moderately Diff. |
| 8 | F | 74 | 30.4 | 12.6 | pT3 | Ad | Well Diff. |
| 9 | M | 79 | 23.5 | 7.6 | pT3 | Ad | Moderately Diff. |

Ad: Adenocarcinoma.

### Excision of *Sptlc2* from mouse intestines decreases ceramides and other sphingolipid biosynthetic intermediates within the intestinal crypts

To better understand the role of sphingolipids in the gut, we produced conditional knockout mice that allow for acute depletion of the *Sptlc2* subunit of SPT from intestinal epithelial cells (IECs). These *Sptlc2*^*δIEC*^ mice were generated by breeding *Sptlc2*^*fl/fl*^ mice that we described previously (Chaurasia et al., 2016) with mice expressing a tamoxifen inducible Cre-recombinase controlled by the villin promoter (Tg(Vil-cre/ERT2)(Goh et al., 2015). Within two days of tamoxifen administration, adult *Sptlc2*^*δIEC*^ mice displayed a marked reduction in *Sptlc2* mRNA expression in the small intestine and colon, as compared to tamoxifen-treated *Sptlc2*^*fl/fl*^ littermates (Figure 2A). The depletion of *Sptlc2* reduced levels of the sphingolipids in the *de novo* synthesis pathway (i.e. ceramides, dihydroceramides, sphinganine, and phytoceramides) within the intestinal crypts (Figure S2A). Surprisingly, it did not alter levels of complex sphingolipids (i.e. sphingomyelins and glucosylceramides) (Figure S2A). The effects of *Sptlc2* depletion manifest predominantly in the crypts, and not the villi; tamoxifen-treatment only affected dihydroceramides, phytoceramides and sphinganine, but not other sphingolipids, within villi of the *Sptlc2*^*δIEC*^ mice (Figure S2B).

**Fig. 2.**
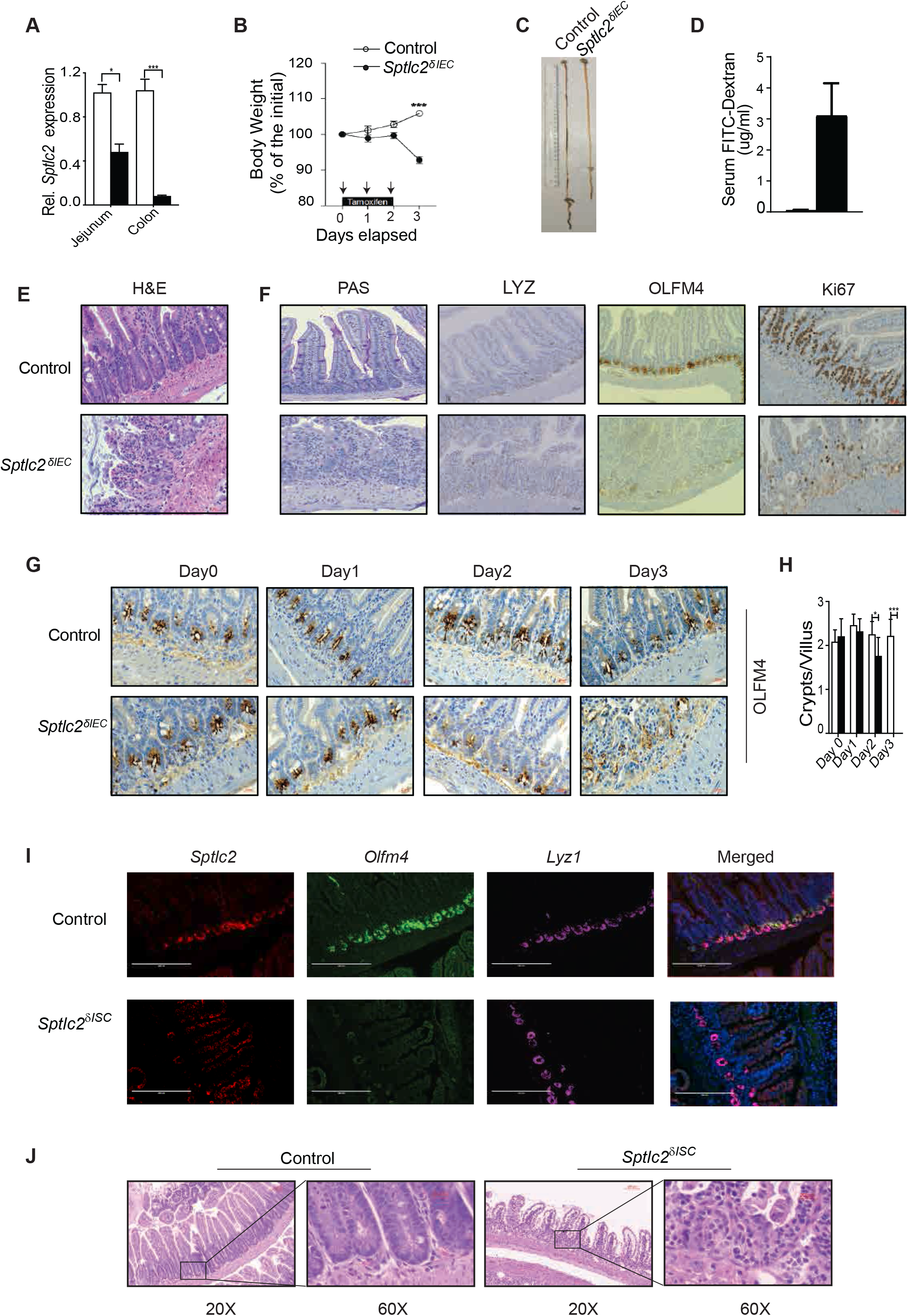
Deletion of *Sptlc2* from intestines disrupts the epithelium and depletes intestinal stem cells. *[A-H] Sptlc2*^*δIEC*^ *and Sptlc2*^*fl/fl*^ *mice were given tamoxifen by intraperitoneal injection (1mg/day for 3 consecutive days).* (A) Measurement of *Sptcl2* levels in jejunum and colon 2-days after they received the first tamoxifen dose. (B) Body weight of *Sptlc2*^*δIEC*^ and *Sptlc*^*fl/fl*^ littermates assessed daily after tamoxifen injection. (C) Image of small intestine collected 4 days after receiving the first tamoxifen dose. (D) Two days after administering the first tamoxifen dose, FITC-dextran was given by oral gavage. Shown are serum levels 4-hours later. (E) H&E staining of the jejunum 4-days after tamoxifen administration. (F) Histology of the jejunum 4-days after tamoxifen including periodic acid–Schiff (PAS) staining for goblet cells, LYZ1 immunostaining for Paneth cells, OLFM4 staining for ISC, and Ki67 staining to identify proliferating cells. (G) Time course study showing OLFM4 positive cells in tissues collected each day after tamoxifen administration. (H) Quantification of crypts per villus from *Sptlc2*^*δIEC*^ and *Sptlc*^*fl/fl*^ mice. *[I-J] Sptlc2*^*δISC*^ *and Sptlc2*^*fl/fl*^ *mice were given tamoxifen by intraperitoneal injection (3mg/day for 5 consecutive days).* (I) *In situ* hybridization (RNAscope) measuring *Sptlc2*, *Olfm4*, and *Lyz1* mRNA expression in the intestine 5 days after tamoxifen injection. Nuclei were stained by DAPI. (J) H&E staining of epithelium from jejunum 5 days after the first tamoxifen dose. Abbreviations: IEC, intestinal epithelia cell; PC, phosphatidylcholine; FITC-Dextran, Fluorescein isothiocyanate– dextran; H&E, Hematoxylin and Eosin staining; PAS, periodic acid-Schiff; *Lyz1*, Lysozyme C1; *Olfm4*, Olfactomedin 4; ISC, intestinal stem cell; serine palmitoyltransferase, long chain base subunit 2; DAPI, 4’,6-diamidino-2-phenylindole.

### Excision of *Sptlc2* from mouse intestines disrupts gut architecture and depletes proliferating ISCs

Deletion of *Sptlc2* led to rapid weight loss (Figure 2B) and death of all animals within 4 days (Figure S2F). Upon necropsy, the *Sptlc2*^*δIEC*^ mice were found to have a short small intestine (Figure 2C and Figure S2C) and a small spleen (Figure S2D). The intestine exhibited increased permeability (Figure 2D), inflammation (Figure S2E). Treatment with broad-spectrum antibiotics prolonged animal survival (Figure S2F), suggesting that sepsis resulting from loss of the intestinal barrier was the ultimate cause of death of the animals. Careful histological assessment revealed severe disruption of the small intestine characterized by disorganized villi and nearly complete loss of crypts (Figure 2E). Following the four-day tamoxifen regimen, the *Sptlc2*^*δIEC*^ mice showed alterations in many cell-types within the villus structure; Goblet cells (PAS), Paneth cells (LYZ1) and ISCs (OLFM4) were depleted and/or mis-localized (Figure 2F). Immunohistochemistry with anti-Ki67 antibodies also revealed a stark decrease in the rapidly proliferating cells located within the crypt (Figure 2F).

To identify which cell types were first affected by *Sptlc2* depletion, we conducted a temporal assessment of the changing pathology, collecting the jejunum through each of the first three days following tamoxifen administration. Though the gross epithelial structure of small intestine in the *Sptlc2*^*δIEC*^ mice remained relatively intact through the first three treatment days, the ISC pool started to disappear within 2-days of tamoxifen injection. This finding was evidenced by reduced expression of OLFM4 (Figure 2G), a disproportionate reduction in the number of crypts (Figure 2H), and fewer proliferating cells in the crypts as assessed by Ki67 staining (Figure S2G and S2H).

### Excision of Sptlc2 from ISCs recapitulates the lethal phenotype of the Sptlc^δIEC^ mice

The findings presented thus far suggest that sphingolipids might be autonomous and essential signals that control the proliferation of ISCs in order to promote epithelial regeneration. Experiments using RNA *in situ* hybridization reinforced this idea, as they revealed that *Sptlc2* transcript levels were much higher in the crypts than they were in the villi (Figure 2I). This observation strongly suggests that the ISCs within the crypt were the major site of sphingolipid synthesis and action. This conclusion was corroborated by the lipidomic assessments showing higher ceramide and phytoceramide levels in the crypts, as compared to the villi (Figure S3A). To selectively study the role of ISC sphingolipids, we generated a mouse line allowing for ISC-specific *Sptlc2* depletion (*Sptlc2^δISC^*) by breeding the aforementioned *Sptlc2*^*fl/fl*^ mice with *Olfm4-EGFP-ires-CreERT2* mice (Schuijers et al., 2014). Injecting tamoxifen into the *Sptlc2*^*δISC*^ mice led to depletion of *Sptlc2* from the crypts, while it was retained in the other cells in the villi (Figure 2I). This led to quick depletion of the ISCs, as assessed by labeling *Olfm4* using RNA *in situ* hybridization (Figure 2I). By comparison, *Lyz1*, which demarcates Paneth cells in the crypt, was largely unaffected by *Sptlc2* depletion (Figure 2I). In these *Sptlc2*^*δISC*^ mice, tamoxifen also recapitulated the gross phenotype of the *Sptlc2*^*δIEC*^ mice, including the reduction in body weight, rapid death, short small intestine and small spleen (Figure S3B-S3F). Histologic analysis showed a comparable disruption of the small intestine structure that was most pronounced in the crypts (Figure 2J).

### Restoring sphingolipids enhances stemness and promotes intestinal organoid survival

To better understand how SPT regulates stem cells, we turned to an intestinal organoid 3-D culture system (Sato et al., 2009). These mini-intestines contain all of the cell types of the mature epithelium but allow for precise control of the tissue environment. Pharmacological inhibition of SPT (i.e. treatment with the SPT inhibitor myriocin) or genetic ablation of *Sptlc2* [i.e. treatment with 4-hydroxytamoxifen (4OHT)] led to rapid death of the organoids, as assessed by propidium iodide staining (Figure 3A). Thus, we recapitulated the *in vivo* phenotype caused by *Sptlc2* depletion using this *in vitro* system. Remarkably, viability of the organoids could be restored by supplementing with short-chain analogs of ceramide (C_2_-ceramide, 25 μM) (Figure 3A). Mass spectroscopy revealed that C_2_-ceramide restored levels of many sphingolipids in either the myriocin or 4OHT treated organoids (e.g. ceramides, glucosylceramides, sphingomyelins and sphingosine), excepting those sphingolipids that cannot be re-produced from C_2_-ceramide via the salvage pathway (e.g. sphinganine and dihydroceramides) (Figure S4A and S4B). In 4OHT-treated organoids, removing C_2_-ceramide from the media re-initiated the death program (Figure 3A).

**Fig. 3.**
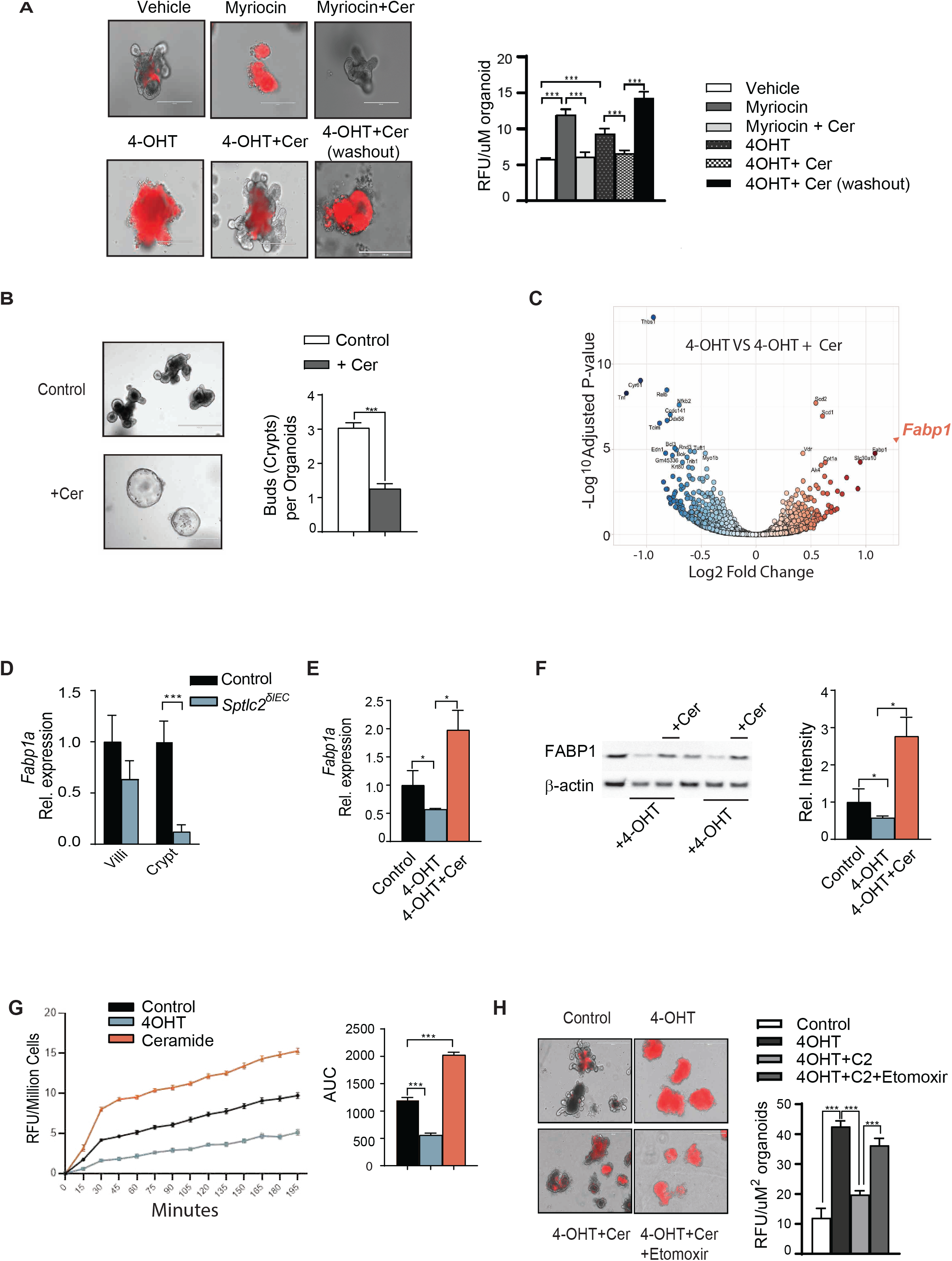
Ceramides promote stemness and intestinal organoid survival by stimulating FABP1 expression to enhance fatty acid uptake. Intestinal organoids were cultured from crypts isolated from the jejunum of *Sptlc*^*δIEC*^ mice. (A) (*left*) Propidium iodide (PI) staining of organoids treated with either vehicle, myriocin (10μM), or 4OHT (200ng/ml) for 48 hours. Some samples were supplemented with C_2_-ceramide (25μM, Cer). In the sample on the lower right, the added C_2_-ceramide was then removed from the media (washout) for the final 24 hours. *(right)* Quantification of the organoid staining reported as relative fluorescence units (RFU) normalized to the area of the organoid. (B) Secondary organoids treated with either vehicle or C_2_-ceramide (25μM) for 7 days. On the right is a quantification of the crypts growing from each organoid. (n=50, *p<0.05, **p<0.01, ***p<0.001). (C) Volcano plot depicting the RNAseq datasets obtained from 4OHT-treated organoids (200 ng/ml) supplemented with or without C_2_-ceramide (25μM) for 48 hours. (D) qPCR analysis of *Fabp1* expression in the villi or crypts from the jejuna of the *Sptlc*^*fl/fl*^ or *Sptlc*^*δIEC*^ mice. (E) qPCR analysis of *Fabp1* gene expression in organoids treated with vehicle, 4OHT (200 ng/ml), or 4OHT+C_2_-ceramide (Cer, 25 μM) for 48 hours. (F) Western blot showing FABP1 and actin expression following *Sptlc2* deletion with 4-OHT and/or C_2_-ceramide supplementation for 48 hours (quantification on the right). (G) Fatty acid uptake into cells dissociated from organoids and treated with vehicle, 4OHT, and/or C_2_-ceramide for 48 hours. The bar graph depicts the area under the curve. (H) *(left)* PI staining images of organoid treated with vehicle, 4OHT, 4OHT plus C2-ceramide or 4OHT plus C2 ceramide plus etomoxir (100μM). *(right)* Quantification of PI staining analyzed by imageJ software (n=30, *p<0.05, **p<0.01, ***p<0.001). Abbreviations: Cer, (d18:1/2:0) N-acetoyl-D-*erythro*-sphingosine; 4-OHT, 4-hydroxytamoxifen; RFU, relative fluorescence unit; IEC, intestinal epithelia cell; FABP1, fatty acid binding protein 1; AUC, area under the curve; Sptlc2, serine palmitoyltransferase; Cpt1a, Carnitine palmitoyltransferase 1A.

To clarify which sphingolipid species might be important for organoid viability, we also exposed these organoid cultures to either the glucosylceramide synthase inhibitor D-threo-et-P4 or the sphingomyelin synthase inhibitor D609, which inhibit the production of glucosylceramides and sphingomyelin, respectively. Neither of these compounds recapitulated the myriocin effects on organoid survival (Figure S4C). Collectively, these data suggest that intermediates in the *de novo* sphingolipid synthesis pathway (e.g. sphingosine, ceramides or phytoceramides), rather than the abundant complex sphingolipids that comprise the majority of the sphingolipidome, (e.g. sphingomyelins), were requisite for organoid viability.

To confirm that C_2_-ceramide modulated stemness, we quantified the number of crypt domains per organoid following exposure to C_2_-ceramide. As we anticipated, treating the secondary organoids with C_2_-ceramide decreased the number of buds sprouting from each organoid, supporting the idea that sphingolipids enhanced stemness (Figure 3B).

### Ceramides enhance stemness by stimulating FABP1 expression to increase fatty acid uptake

To explore the molecular mechanism(s) that might account for the profound sphingolipid actions in the intestinal epithelium, we conducted RNA sequencing on *Sptlc2*-knockout organoids treated with or without C_2_-ceramide. *Fabp1*, which encodes a fatty acid binding protein (i.e. FABP1) involved in lipid uptake, was amongst the handful of transcripts most dramatically upregulated by C_2_-ceramide (Figure 3C). This C_2_-ceramide effect on FABP1 was particularly interesting owing to a bevy of studies showing that saturated fatty acids increase the expression of FABP1 in ISCs (Mihaylova et al., 2018) and that increasing fatty acid oxidation enhances stemness (Bensard et al., 2019; Knobloch et al., 2017; Mihaylova et al., 2018). *Sptlc2* depletion from *Sptlc2*^*δIEC*^ mice decreased *Fabp1* transcripts in the crypt, but not the villi (Figure 3D). Using organoids, we also confirmed that *Sptlc2* depletion decreased, and C_2_-ceramide restored, expression of *Fabp1* mRNA and FABP1 protein (Figure 3E and 3F).

To evaluate the functional consequences of these interventions on lipid handling, we quantified fatty acid uptake in organoids isolated from the *Sptlc2*^*δIEC*^ mice treated with vehicle, 4OHT, or C_2_-ceramide. C_2_-ceramide stimulated fatty acid uptake in organoids. By contrast, depletion of *Sptlc2* using 4OHT reduced rates of fatty acid uptake (Figure 3G).

The RNA sequencing also identified the transcript encoding carnitine palmitoyltransferase-1 (*Cpt1a*), the enzyme that facilitates fatty acid entry into mitochondria, as a C_2_-ceramide-responsive gene. We confirmed that C_2_-ceramide slightly increased expression of *Cpt1a* transcripts (by qPCR, Figure 4A) and CPT1a protein (by Western blot, Figure 4B). The CPT1 inhibitor etomoxir negated the protective actions of C_2_-ceramide on survival of *Sptlc2* knockout organoids (Figure 3H). Thus, induction of fatty acid uptake and utilization were essential C_2_-ceramide actions that preserved ISC viability.

**Fig. 4.**
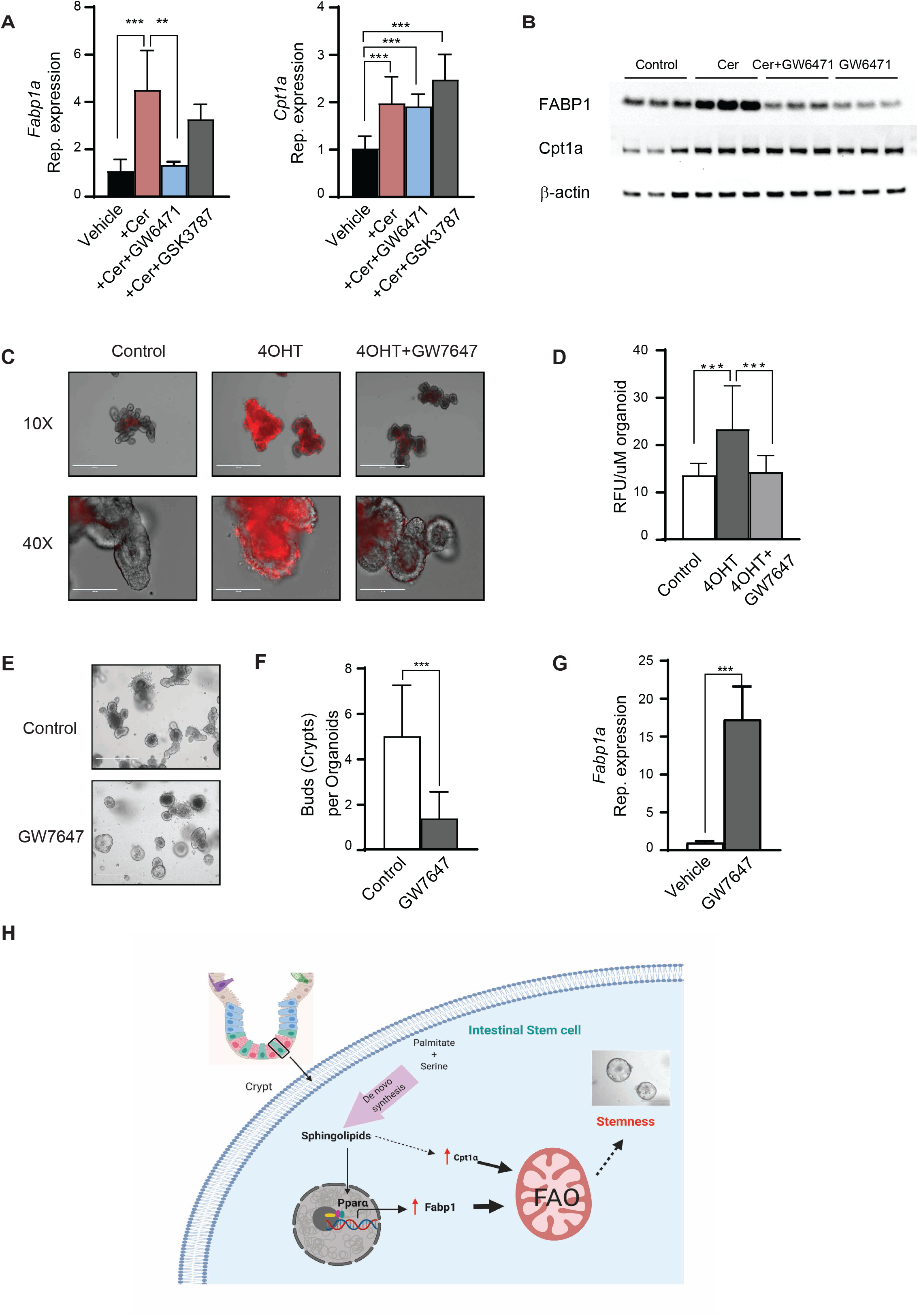
Ceramides increases FABP1 expression through PPARα activation. (A) qPCR analysis of *Fabp1* and *Cpt1a* expression in organoids treated with vehicle, C2-ceramide (25 μM), and C_2_-ceramide with GW6471(PPARα antagonist, 1μM) or GSK3787 (PPARδ antagonist, 1μM) for 48 hours. (B) Western blot analysis of FABP1, CPT1a, and actin in organoids treated with C_2_-ceramide with or without the PPARα antagonist GW6471(GW6471, 1μM) for 48 hours. (C) Propidium iodide (PI) staining of organoids treated with either vehicle, 4OHT (200ng/ml) or 4OHT with the PPARα agonist GW7647 for 48 hours. (D) Quantification of PI staining of 4C. (E) Secondary organoids treated with either vehicle or GW7647 (1μM) for 7 days. (F) Quantification of the crypts growing from each organoid. (n=50, *p<0.05, **p<0.01, ***p<0.001). (G) qPCR analysis of *Fabp1* expression in organoids treated with either vehicle or GW7647 (1μM). (H) Schematic depicting the concept shown herein that SPT-generated sphingolipids are obligate intermediates linking saturated fatty acids and serine to stemness. The picture is generated using Biorender software (Biorender.com). Abbreviations: Cer, (d18:1/2:0) N-acetoyl-D-*erythro*-sphingosine; 4-OHT, 4-hydroxytamoxifen; RFU, relative fluorescence unit; IEC, intestinal epithelia cell; FABP1, fatty acid binding protein 1; Cpt1a, Carnitine palmitoyltransferase 1A; PPARα, Peroxisome proliferator-activated receptor alpha; AUC, area under the curve; FAO, fatty acid oxidation.

### Ceramides increase FABP1 expression through PPARα activation

Prior studies have shown that the transcription factors peroxisome-proliferator activated receptor isoforms alpha (PPARα) and delta (PPARδ) regulate FABP1 expression in the small intestine (Darimont et al., 1998; Hughes et al., 2015; Mochizuki et al., 2001; Poirier et al., 2001). In addition, recent studies have implicated PPARδ as a critical intermediate linking saturated fats to the control of fatty acid oxidation and the enhancement of ISC stemness (Beyaz et al., 2016; Mihaylova et al., 2018). We thus tested whether *Fabp1* or *Cpt1* expression was controlled by either of these transcription factors. The PPARα antagonist GW6471 blocked the C_2_-ceramide induction of *Fabp1* (Figure 4A and 4B), but not *Cpt1a*. The PPARδ antagonist GSK 3787 had no effect on either gene (Figure 4A). Thus, PPARα, but not PPARδ, was an essential intermediate linking C_2_-ceramide to the induction of FABP1.

Lastly, we evaluated whether PPARα was sufficient to maintain organoid viability, even in the absence of *Sptlc2*. The *PPARα* agonist GW7647 restored viability of organoids lacking *Sptlc2* (Figure 4C and 4D). GW7647-treated organoids had fewer crypts in each organoid, thus recapitulating the C_2_-ceramide effect on organoid stemness (Figure 4E and 4F). Moreover, PPARα activation recapitulated the ceramide effect on *Fabp1* expression (Figure 4G). These studies convincingly display the existence of a sphingolipid-PPARα-FABP1 axis that links macronutrients to the maintenance of the ISC pool.

## Discussion

These studies provide important mechanistic insight about the nutritional signals that control the metabolism and proliferation of ISCs, the cell-of-origin of gastrointestinal cancers. Prior studies have shown that the transition from normal ISCs into a hyperplastic lesion is dependent upon fuel choice (Bensard et al., 2020). Excessive fatty acid supply increases the proliferation and stemness of ISC and progenitors within the crypt to promote tumorigenicity (Bensard et al., 2019; Knobloch et al., 2017; Mihaylova et al., 2018). In mouse models of colorectal cancer, genetic depletion or pharmacological inhibition of the mitochondrial pyruvate carrier, which increases dependence on fatty acids oxidation, expands the stem cell compartment and doubles the frequency of adenoma formation (Bensard et al., 2020; Schell et al., 2017). By contrast, germline deletion of *Fabp1* slows lipid uptake and reduces adenoma formation (Dharmarajan et al., 2013). The findings described herein reveal that sphingolipids produced from ectopic palmitate and serine influence this fuel choice by enhancing fatty acid import through FABP1.

Our data further identify PPARα as an essential intermediate that links endogenous sphingolipids to the induction of *Fabp1*. This is consistent with prior studies, which show that high fat diet increases levels of *Pparα* and its downstream target *Fabp1* in rat jejunum (Suruga et al., 1999), as well as those showing that a PPARα antagonist induces organoid death (Stine et al., 2019). Moreover, it aligns with studies showing that sphingolipids increase PPARα-mediated transcription (Correnti et al., 2018; Murakami et al., 2011; Tsuji et al., 2009; Van Veldhoven et al., 2000). Ceramides, phytoceramides, and sphingoid bases have been identified as upstream activators of PPARα, with some synthetic sphingolipids showing direct binding capabilities (Murakami et al., 2011; Tsuji et al., 2009; Van Veldhoven et al., 2000). Our findings reveal that one or more of these sphingolipids serves as a signal to increase fatty acid uptake and oxidation, and thus provides important new information about the nutritional sensing machinery that induces this transcriptional program.

The studies are also important for discerning the precise role of SPT in the gut. Prior studies have shown that genetic ablation of *Sptlc2* (Li et al., 2018; Ohta et al., 2009) or pharmacological inhibition of SPT (Genin et al., 2016) has deleterious consequences on gut health, with the former causing animal death and the latter gastric enteropathy. Indeed, inflammatory bowel disease has been associated with decreased expression of SPT (Li et al., 2018). The studies described herein indicate that these pathogenic states result from decreased conversion of free fatty acid and amino acids into ceramides and a concomitant depletion of the ISC pool. Thus, strategies targeting sphingolipids could prove effective as a means of modulating the health of the intestinal barrier.

Collectively, these data support the conclusion that overproduction of sphingolipids enhances stemness of cells within the intestinal crypts. Using the organoid system, we identified an intriguing mechanism linking sphingolipids to the modulation of stemness through their effects on PPARα-mediated induction of FABP1 and a resultant increase in fatty acid uptake. These observations help reveal how saturated fatty acids control intestinal stemness and provide an explanation for the utility of serine deprivation as a means to decrease tumor incidence. Finally, because stem cells play a major role not only in tissue regeneration but also in carcinogenesis, deciphering their response to dietary factors is of the utmost importance for the treatment of degenerative conditions. Ultimately, these studies could suggest dietary or pharmacological strategies for maximizing regenerative capacity while also minimizing the risk of cancer development.

## Author contributions

S.A.S, B.C., and Y.L. conceived of the project designed the experiments and wrote the manuscript. Y.L, B.C, V.A.K., J.A.M., performed experiments and analyzed data. C.B, J.R, H.C, Y.H provided some of the mouse strains used in this study. J.L.W and W.L.H. assisted in quantification of images. D.L provided human tissues used in this study and J.C, P.W and P.J.M aided in lipid analysis.

## Acknowledgements

We wish to acknowledge the support from Metabolomics, Histology, and Metabolic Phenotyping Cores at the Health Sciences Center of the University of Utah. All data are available in the main text or the supplementary materials.

## Materials and Methods

### Human colon samples

Human colon cancer tissues were obtained from the Preclinical Research Resource at the Huntsman Cancer Institute. Tumors were collected during tumor resection (not biopsies) and cryopreserved as viable tissue. The adjacent normal tissue was collected as a paired control.

### Animal Colonies

All animal experiments were conducted with protocols approved by the Institutional Animal Care and Use Committee (IAVUC) of the University of Utah. <u>Generation of *Sptlc2*^*δIEC*^ *Mice*</u>. *Sptlc2*^*fl/fl*^ mice were crossed with Villin-Cre-ERT2 mice(Goh et al., 2015) to generate gut-specific *Sptlc2* knockout animals. To deplete the gene, mice were injected intraperitoneally with tamoxifen (1mg/mouse) every day for three days at 8-10 weeks of age. *Sptlc2*^*fl/fl*^ mice littermates that were also injected with tamoxifen were used as controls. <u>Generation of *Sptlc2*^*δISC*^ *Mice*</u>. *Sptlc2*^*fl/fl*^ mice were crossed with Olfm4-EGFP-ires-CreERT2 mice *(26)* to generate ISC-specific *Sptlc2* knockout animals. To deplete the gene, mice were injected intraperitoneally with tamoxifen (3mg/mouse) every day for five days at 8-10 weeks of age. *Sptlc2*^*fl/fl*^ mice littermates that were also injected with tamoxifen were used as controls.

### Sphingolipidomics

#### University of Utah Metabolomics Core

Tissues or organoids suspended in 120 ◻1 PBS went through three freeze-thaw cycles to homogenize the samples. Thereafter, 100 ◻1 of homogenate was used for lipid extraction and 20 ◻1 was saved for protein assays. Samples were spiked with 225 μL MeOH containing internal standards (deuterated ceramide mixture (C16, 60 pmol/sample; C18, 35 pmol/sample; C24, 150 pmol/sample; C24:1, 312 pmol/sample); dihydroceramide C18:1, 5pmol/sample; deuterated sphingomyelin, 50 pmol/sample; glucosylceramide C17, 50 pmol/sample; and deuterated S1P, 5pmol/sample) followed by 750 μL MTBE (methyl *tert*-butyl ether). The samples were then incubated for 60 min on ice with brief vortexing every 15 min. Following centrifugation for phase separation, the upper phase was transferred to another Eppendorf tube which was dried using a Speedvac. The dried, extracted lipid was reconstituted in methanol:toluene (9:1). The lipid extracts were then analyzed by liquid chromatography-mass spectrometry (LC/MS). Briefly, lipid extracts were separated on an Acquity UPLC CSH C18 1.7 μm 2.1 × 50 mm column maintained at 60 °C connected to an Agilent HiP 1290 Sampler, Agilent 1290 Infinity pump, and Agilent 6490 triple quadrupole (QqQ) mass spectrometer. Sphingolipids were detected using dynamic multiple reaction monitoring (dMRM) in positive ion mode. Source gas temperature was set to 210°C, with a gas (N_2_) flow of 11 L/min and a nebulizer pressure of 30 psi. Sheath gas temperature was 400°C, sheath gas (N_2_) flow of 12 L/min, capillary voltage is 4000 V, nozzle voltage 500 V, high pressure RF 190 V and low-pressure RF is 120 V. Injection volume was 2 μL and the samples were analyzed in a randomized order with the pooled QC sample injection eight times throughout the sample queue. Mobile phase A consisted of ACN:H_2_O (60:40 v/v) and mobile phase B consists of IPA:ACN:H_2_O (90:9:1 v/v) both in 10 mM ammonium formate and 0.1% formic acid. The chromatography gradient started at 15% mobile phase B, increased to 30% B over 1 min, increased to 60% B from 1-2 min, increased to 80% B from 2-10 min, and increased to 99% B from 10-10.2 min where it was held until 14 min. Post-time was 5 min and the flowrate was 0.35 mL/min throughout. Collision energies and cell accelerator voltages were optimized using sphingolipid standards with dMRM transitions as [M+H]^+^→[*m/z* = 284.3] for dihydroceramides, [M+H]^+^→[*m/z* = 287.3] for isotope labeled dihydroceramides, [M-H_2_O+H]+→[*m/z* = 264.2] for ceramides, [M-H_2_O+H]^+^→[*m/z* = 267.2] for isotope labeled ceramides and [M+H]^+^→[M-H_2_O+H]_+_ for all targets. Sphingolipids without available standards were identified based on HR-LC/MS, quasi-molecular ion and characteristic product ions. Their retention times were either taken from HR-LC/MS data or inferred from the available sphingolipid standards. Results from LC/MS experiments are collected using Agilent Mass Hunter Workstation and analyzed using the software package Agilent Mass Hunter Quant B.07.00. Sphingolipids were quantitated based on peak area ratios to the standards added to the extracts.

#### Baker Heart and Diabetes Institute Metabolomics Laboratory

The quantification of specific lipids at the Baker Heart and Diabetes Institute Metabolomics laboratory was done as follows: 200 μL of chloroform/methanol (2:1, v/v) were added together with 10 μL ISTD in chloroform/methanol (1:1, v/v) to 25-60μg of protein lysate in 10μl cold PBS. The mixture was mixed for 10 min on a rotary mixer, sonicated in a water bath (18 °C–24 °C) for 30 min, left to stand on the bench for 20 min and then centrifuged (16,000 × g, 10 min, 20 °C). The supernatant was transferred to a 96-well plate and dried under a stream of nitrogen gas at 40 °C. Samples were reconstituted with 50 μL H_2_O-saturated 1-butanol and sonicated for 10 min. Finally, 50 μL of 10 mM ammonium formate in MeOH was added. The extract was centrifuged (1700 × g, 5 min, 20 °C) and the supernatant was transferred into a 0.2 mL glass insert with Teflon insert cap for analysis by LC ESI-MS/MS as described *(27)*.

### Isolation, culture, and analysis of intestinal organoids

Small intestinal crypts were isolated and grown into organoids as described previously(Sato et al., 2009). Isolated small intestines were chopped into small pieces and washed in cold PBS. They were then incubated in 2 mM EDTA with PBS for 30 min on ice. EDTA medium was removed and the tissue fragments were washed by cold PBS. They were then pipetted vigorously to dissociate and enrich the crypts into the supernatant. Isolated crypts were pull down by centrifuged at 200*g*, and then re-suspended in Matrigel (Car#354277, BD) for organoids culture. Myriocin (10uM) was used to inhibit SPT2 activity in the organoids for 24-48 hours. Deletion of *Sptlc2* in organoid cultures from *Sptlc2*^*μIEC*^ mice was induced by addition of 200ng/ml 4-OH-tamoxifen (4-OHT) to the culture medium for 48 hours. Cell death of organoid cultures was determined by propidium iodine (PI)-positive staining. Images were taken by EVOS digital inverted microscope. Quantification of PI staining was done by Image J software (NIH).

### Antibiotic treatment of *Sptlc2*^*δIEC*^ mice

A broad-spectrum antibiotic mixture was added to the drinking water one month before the ablation of the *Sptlc2* gene(Takahashi et al., 2014). The cocktail included 200mg ciprofloxacin (Sigma-Aldrich), 1g ampicillin (Sigma-Aldrich), 100mg metronidazole (Sigma-Aldrich), 500mg vancomycin (Labconsult), 1g neomycin, 2.50g streptomycin (ThermoFisher), 100,000U penicillin (ThermoFisher), 1g bacitracin (Sigma-Aldrich) and 1g ceftazidime (Sigma-Aldrich) were dissolved in 1 liter of water. Feces was collected and cultured in liquid cultures of Thioglycollate medium (Sigma) and McConkey agar plates. No bacteria growth was observed from both cultures.

### Quantitative RT-PCR

Total RNA was extracted from tissues or cells using the RNeasy Mini Isolation Kits with DNA digestion (QIAGEN) according to the manufacturer’s instructions. Isolated total RNA was reverse-transcribed into cDNA using commercially available kits (Quantbio). All subsequent qRT-PCR reactions were performed on a QuantStudio 12K Flex Real-Time PCR system (Thermo Fisher Scientific) using the Qiagen QuantiFast SYBR Green PCR kit. qPCR primers were purchased from Sigma-Aldrich, KiCqStart Primers. For normalization threshold cycles (C_t_-values) of all replicate analyses were normalized to *Ppib* or B2M for mouse tissues and human tissue within each sample to obtain sample-specific ΔC_t_ values (= C_t_ gene of interest - C_t_ *Ppib*, B2M). To compare the effect of various treatments with untreated controls, 2^−ΔΔCt^ values were calculated to obtain fold expression levels, where ΔΔC_t_ = (ΔC_t_ treatment - ΔC_t_ control).

### FITC-Dextran Permeability Test

Intestinal permeability was assessed by oral gavage of FITC–dextran (Sigma-Aldrich). Mice were administered 500◻ FITC-Dextran (20mg/ml in PBS, 10mg/mouse) by gavage. Serum sample were collected 4-hours after oral gavage; Fluorescence were measured by a fluorescence plate reader at 488nm. Serum FITC-Dextran concentration was calculated according to the standard curve made by a serial dilution from the stock in PBS.

### RNAseq Analysis

Total RNA was extracted and RNAseq analysis was conducted at University of Utah genomics core facility who evaluated the RNA quality by NanoDrop spectrophotometer and Agilent Bioanalyzer RNA 6000 Chip and further performed the sequencing by Illumina HiSeq2000 with 50 cycle paired end sequencing. Data were analyzed by bioinformatics analysis service at the microarray core facility at the University of Utah. The criteria for differential expression included only genes with FDR of <0.05 or log-transformed false discovery rate (FDR) of >13 and with a fold change greater than or equal to ±1.5 relative to controls. The Fragments Per Kilobase Million (FPKM) values of the differentially expressed genes were log transformed and normalized to the average of the controls. These values were used to produce hierarchical clustering and generate heat maps of the genes using Cluster 3.0 and Java Tree View open source programs, respectfully.

### Western Blot Analysis

Proteins were extracted from organoids by homogenizing in RIPA buffer (0.5% NP-40, 0.1% sodium deoxycholate, 150 mM NaCl, 50 mM Tris-HCl, pH 7.5) containing protease inhibitors (Complete Mini, Roche). The homogenate was cleared by centrifugation at 4°C for 30 min at 15,000 g and the supernatant containing the protein fraction recovered. Protein concentration in the supernatant was determined using the BCA Protein Assay Kit (Pierce). 10 μg of proteins was resolved using precast Bolt (4-12% Bis-Tris gels, Invitrogen) and transferred to Nitrocellulose membranes (GE Healthcare). Membranes were blocked with 5% BSA in Tris-buffered saline containing 0.2% Tween-20 (TBS-T) and incubated with primary antibodies at 4°C overnight.

### Histology and Immunochemistry

For histology, tissues were fixed in 10% buffered formalin for 48 hours, embedded in paraffin, sectioned at 5μm and mounted on the slides. The slides were either stained with hematoxylin and eosin or PAS for goblet cells. <u>Immunohistochemistry</u>: The slides were sent to Arup laboratories for immunohistochemical analysis of Olfm4, Lyz1, Ki67, F4/80 and cleaved caspase-3 staining.

### Fatty Acid Uptake Assay

Organoids were grown and treated with vehicle, 4OHT (200ng/ml) or C_2_-ceramide for 48 hours. Organoids were then dissociated into single cell suspension and seeded onto a 384-well plate. BODIPY-dodecanoic acid fluorescent fatty acid analog was added (QBT™ Fatty Acid Uptake Assay Kit, Molecular Devices) and the plate was immediately transferred to a fluorescence microplate reader for kinetic reading (every 20 seconds for 30-60 minutes) using a bottom-read mode. Measurement was done with an excitation filter of 480 nm and an emission filter of 520.

### RNA fluorescence in situ hybridization

Formalin-fixed paraffin-embedded (FFPE) mouse jejunum sections were de-paraffinized, rehydrated, and then hybridized with mRNA probes against mouse *Sptlc2, Olfm4* and *Lyz1* in accordance with the manufacturer’s instructions (Advanced Cell Diagnostics, Newark, CA). In brief, FFPE sections were pretreated with hydrogen peroxide, incubated in Target Retrieval solution in a steamer for 30 minutes, and permeabilized by incubating in Protease Plus solution for 40 minutes. After hybridization, a fluorescent kit was used to amplify the mRNA signal. TSA Plus Fluorescein, TSA Plus Cyanine 3, and TSA Plus Cyanine 5 fluorescent signals were detected using EVOS digital inverted microscope.

## Statistical analysis

Data were plotted as the mean ± SEM. Student *t*-test, one-way or two-way ANOVA were carried out using Excel or prism and statistical significance was considered meaningful at p<0.05.

---

**Figure. S1.**
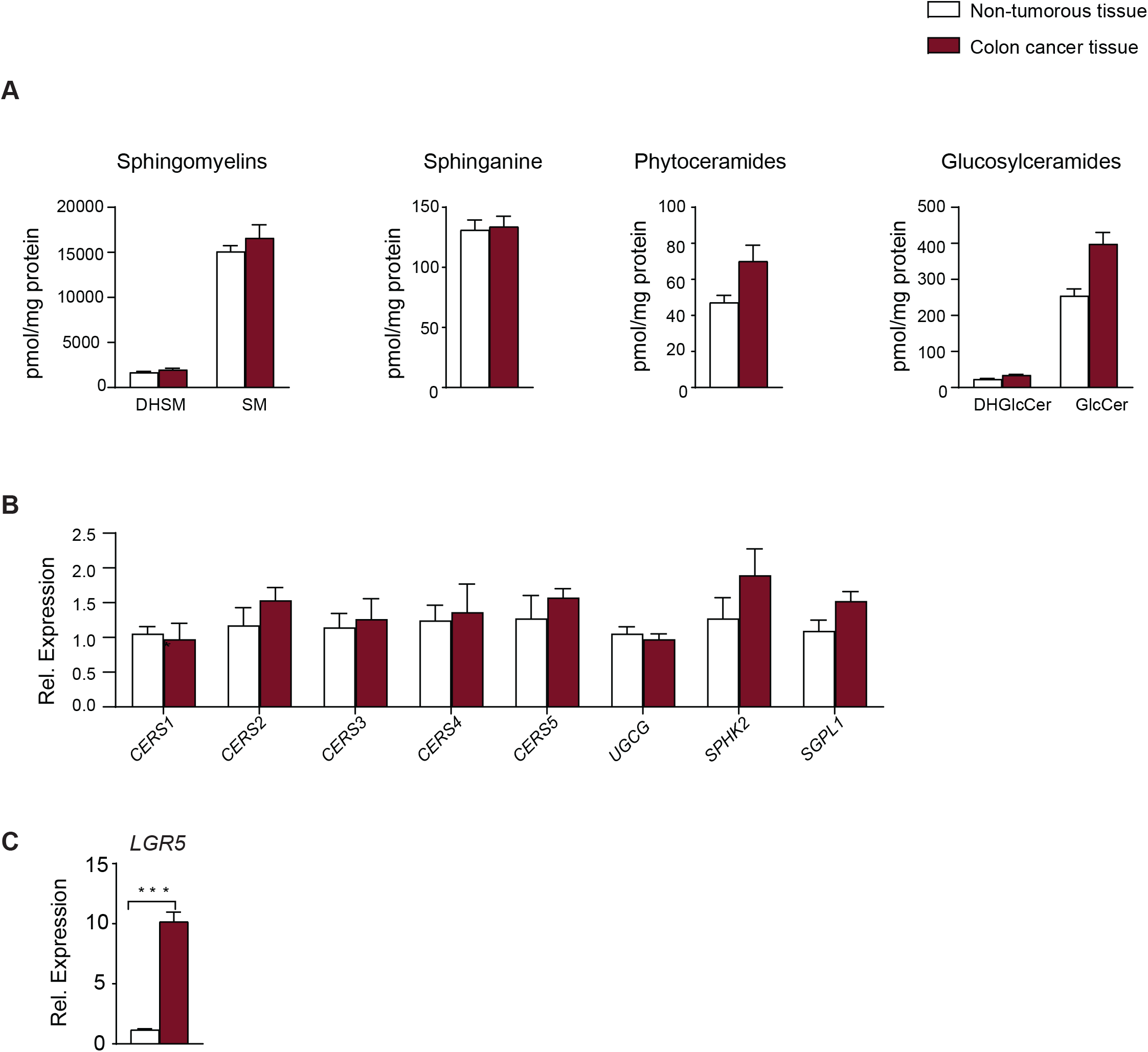
(related to Figure 1). Quantification of Sphingolipids in Colorectal Cancer. Quantification of (A) sphingolipids, (B) transcripts encoding genes involved in sphingolipid metabolism (using qPCR), and (C) transcripts encoding LGR5 in colon cancer tissue and adjacent non-tumorous tissue. (n=9) (*p<0.05, **p<0.01, ***p<0.001, n=9). Abbreviations: DHSM, dihydrosphingomyelin; SM, sphingomyelin; DHGlcCer, dihydroglucosylceramide; GlcCer, glucosylceramide; CERS1, ceramide synthase-1; CERS2, ceramide synthase-2; CERS3, ceramide synthase-3; CERS4, ceramide synthase-4; CERS5, ceramide synthase-5; UGCG, Ceramide glucosyltransferase; SPHK2, sphingosine kinase 2; SGPL1, sphingosine phosphate lyase 1; LGR 5, Leucine-rich repeat-containing G-protein coupled receptor 5.Type or paste caption here. Create a page break and paste in the Figure above the caption.

**Figure. S2.**
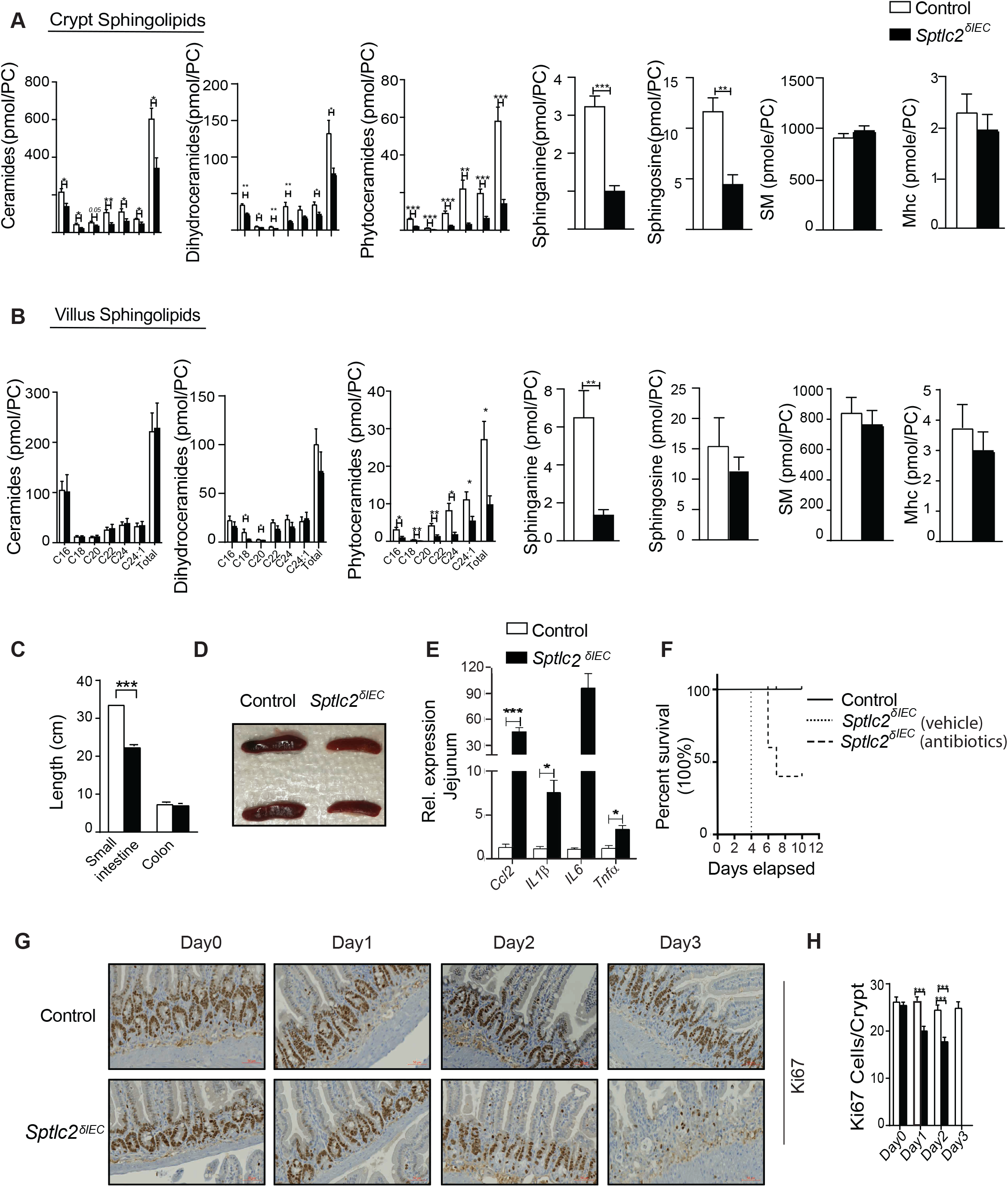
(related to Figure 2).Deletion of *Sptlc2* from intestines disrupts the epithelium and depletes intestinal stem cells. (A) Quantification of sphingolipids in the crypt from the jejunum of *Sptlc2*^*δIEC*^ and *Sptlc*^*fl/fl*^ littermates 2-days after mice received the first tamoxifen dose. (B) Quantification of sphingolipids in villi isolated from the jejuna of Sptlc2δIEC and Sptlcfl/fl littermates two days after tamoxifen treatment. (C) Quantification of small intestine length in SptlcδIEC and Sptlc2fl/fl littermates four days after tamoxifen treatment (n=6). (D) Image of spleens harvested four days after tamoxifen treatment. (E) qPCR analysis of inflammatory markers four days after tamoxifen injection. (F) Survival curve of *Sptlc2*^*δIEC*^ and *Sptlc*^*fl/fl*^ mice treated with or without antibiotics. (G) Time course study showing Ki67 positive cells in the small intestine from *Sptlc2*^*δIEC*^ and *Sptlc*^*fl/fl*^ mice. (H) Quantification of Ki67 positive cells in each crypt. Abbreviations: PC, phosphatidylcholine; IEC, intestinal epithelia cell; SM, sphingomyelin; Mhc, monohexosylceramide; Ccl2, chemokine ligand 2; IL1, interleukin 1-beta; IL6, interleukin 6; Tnf, tumor necrosis factor alpha; H&E, Hematoxylin and Eosin staining.

**Figure. S3.**
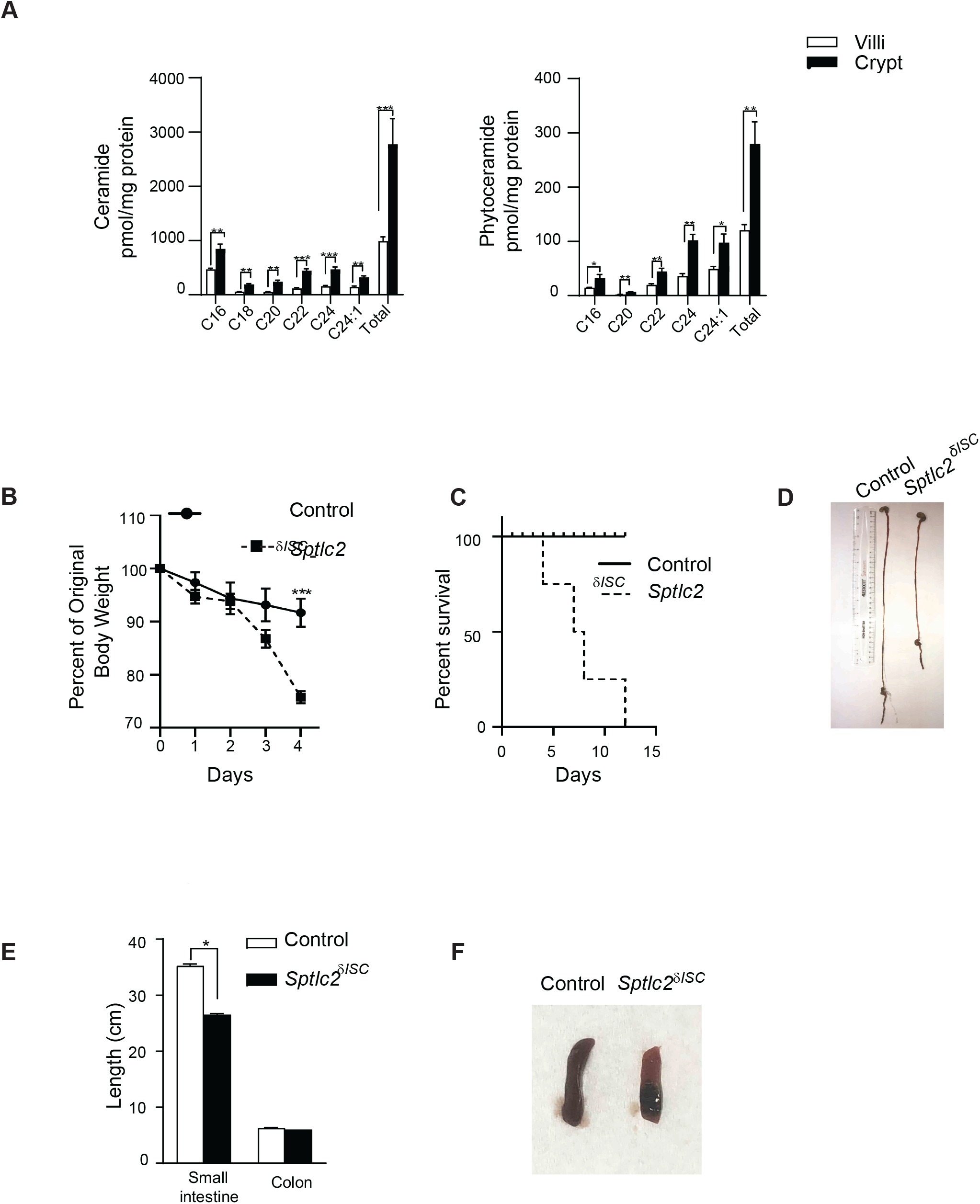
(related to Figure 2). Deletion of Sptlc2 from intestinal stem cells (SptlcδISC) recapitulates the pathologic phenotype of the SptlcδIEC mice. (A) Quantification of sphingolipids in the villi or crypt of small intestine from a wide type C57BL/6 mouse. (B) Body weight of mice after *Sptlc2* gene deletion. (C) Survival curve of *Sptlc2*^*δISC*^ and *Sptlc*^*fl/fl*^ littermates. (D) Image of small intestine after Sptlc2 deletion. (E) Quantification of the length of the intestines. (F) Image of spleens harvested four days after tamoxifen treatment. Abbreviations. ISC, intestinal stem cell; H&E, Hematoxylin and Eosin staining.

**Figure. S4.**
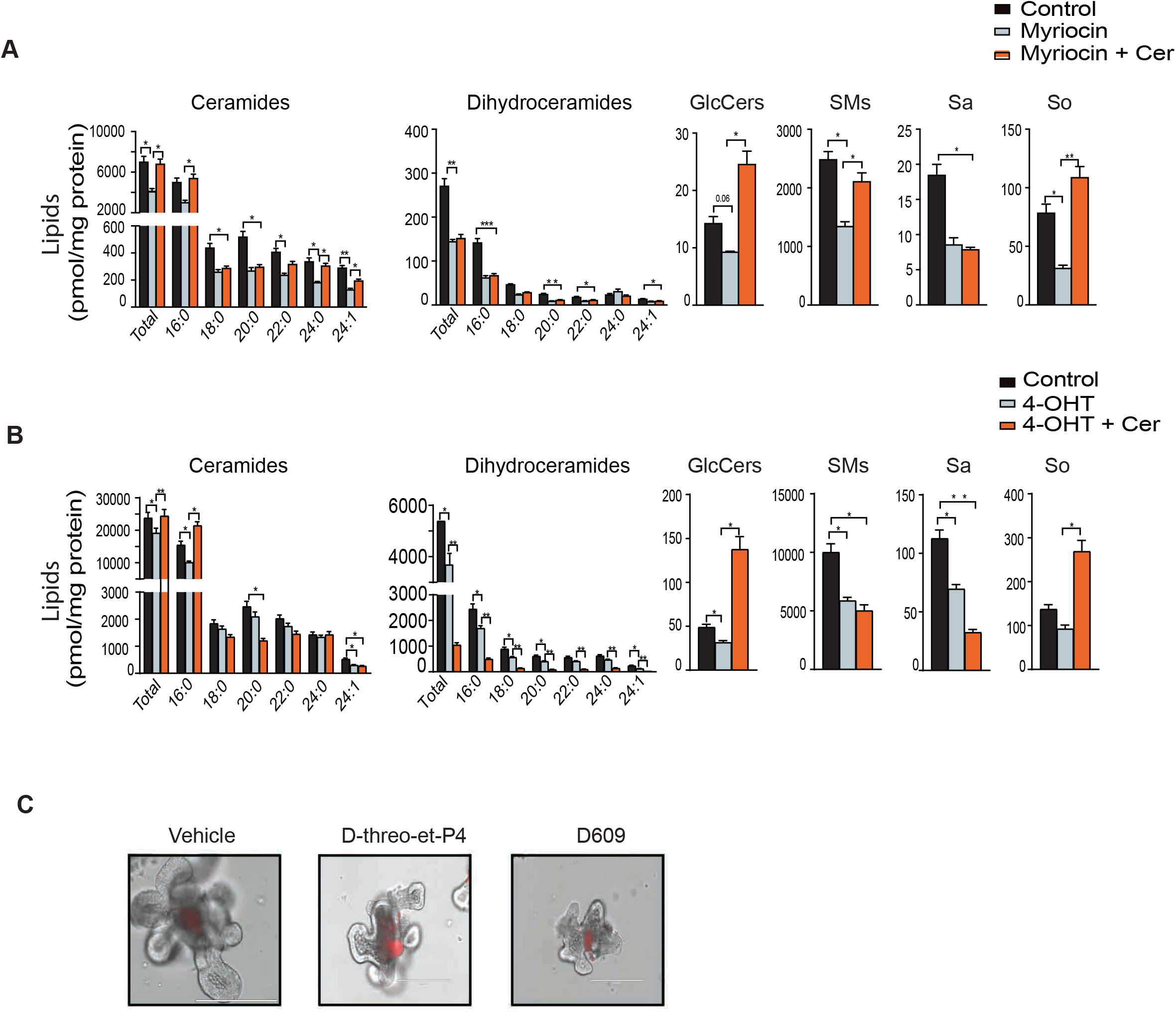
(related to Figure 3 and 4).Quantification of Sphingolipids in Organoids. (A) Quantification of sphingolipids in organoids treated with vehicle, myriocin (10μM), or myriocin plus C2-ceramide (25μM) for 24 hours. (B) Quantification of sphingolipids in organoids treated with vehicle, 4OHT (200ng/ml) or 4OHT plus C2 ceramide (n=6, *p<0.05, **p<0.01, ***p<0.001). (C) PI staining of organoids treated with vehicle, De-threo-et-P4 (100nM, GCS inhibitor) or D609 (100nM, SMS inhibitor) for 72 hours. Abbreviations: GlcCers, glucosylceramides; SMs, sphingomyelins; Sa, sphinganine; So, sphingosine; Cer, Ceramide (d18:1/2:0), N-acetyl-D-erythro-sphingosine; 4OHT, 4-hydroxytamoxifen.

